# Assessing uncertainty in the rooting of the SARS-CoV-2 phylogeny

**DOI:** 10.1101/2020.06.19.160630

**Authors:** Lenore Pipes, Hongru Wang, John P. Huelsenbeck, Rasmus Nielsen

## Abstract

The rooting of the SARS-CoV-2 phylogeny is important for understanding the origin and early spread of the virus. Previously published phylogenies have used different rootings that do not always provide consistent results. We investigate several different strategies for rooting the SARS-CoV-2 tree and provide measures of statistical uncertainty for all methods. We show that methods based on the molecular clock tend to place the root in the B clade, while methods based on outgroup rooting tend to place the root in the A clade. The results from the two approaches are statistically incompatible, possibly as a consequence of deviations from a molecular clock or excess back-mutations. We also show that none of the methods provide strong statistical support for the placement of the root in any particular edge of the tree. Our results suggest that inferences on the origin and early spread of SARS-CoV-2 based on rooted trees should be interpreted with caution.

## Introduction

SARS-CoV-2, the virus causing COVID-19 or ‘Severe Acute Respiratory Syndrome,’ has a single-stranded RNA genome 29,891 nucleotides in length (Wu *et al.*, 2020; Zhou *et al.*, 2020b). The exact origin of the virus causing the human pandemic is unknown, but two coranoaviruses isolated from bats — RaTG13 isolated from *Rhinolophus affinis* (Zhou *et al.*, 2020a) and RmYN02 isolated from *Rhinolophus malayanus* (Zhou *et al.*, 2020b), both from the Yunnan province of China — appear to be closely related. After accounting for recombination, the divergence time between these bat viruses and SARS-CoV-2 is estimated to be approximately 52 years [95% C.I. (28, 75)] and 37 years [95% C.I. (18,56)] (Wang *et al.*, 2020), for RaTG13 and RmYN02 respectively, using a strict clock, only the most closely related sequences, and only synonymous mutations, or 51 years [95% HPD credible interval (40, 70)] for RaTG13 (Boni *et al.*, 2020) using a relaxed clock and all mutations including divergent sequences saturated in synonymous sites. After the emergence of the virus was first reported from Wuhan in China (Li *et al.*, 2020b) it rapidly spread to many other areas of the world (World Health Organization, 2020). However, the events leading to the early spread of the viruses are still unclear, in part because there is substantial uncertainty about the rooting of the SARS-CoV-2 phylogeny. The importance in identifying the origin of the virus has prompted other analyses on the uncertainty of rooting the phylogeny (Gomez-Carballa *et al.*, 2020; Morel *et al.*, 2020). Previous analyses have reached different conclusions about the rooting of the phylogeny. While analyses that used an outgroup reached one placement (Shen *et al.*, 2020; Tang *et al.*, 2020; Yu *et al.*, 2020; Zhang *et al.*, 2020), analyses that used midpoint rooting reached another placement (Li *et al.*, 2020c, d; Nie *et al.*, 2020), and yet other analyses using a molecular clock have reached a different placement of the root (Benvenuto *et al.*, 2020; Giovanetti *et al.*, 2020; Lemey *et al.*, 2020; Li *et al.*, 2020a). In particular, there is considerable discrepancy between rootings based on rooting with the two closest outgroup sequences (Fig 1A), which has a rooting in clade A, and rooting based on a molecular clock (Fig 1B), which has a rooting in clade B, using clade designations by Rambaut *et al.* (2020). Clade B contains the earliest sequences from Wuhan, and a rooting in this clade would be compatible with the epidemiological evidence of an origin of SARS-CoV-2 in or near Wuhan. However, if an outgroup rooting is assumed (Fig 1A) the inferred origin is in Clade A which consists of many individuals from both inside and outside East Asia. Such a rooting would be compatible with origins of SARS-CoV-2 outside of Wuhan. The rooting of the SARS-CoV-2 pandemic is, therefore, critical for our understanding of the origin and early spread of the virus. However, it is not clear how best to root the tree and how much confidence can be placed in any particular rooting of the tree.

**FIG. 1.**
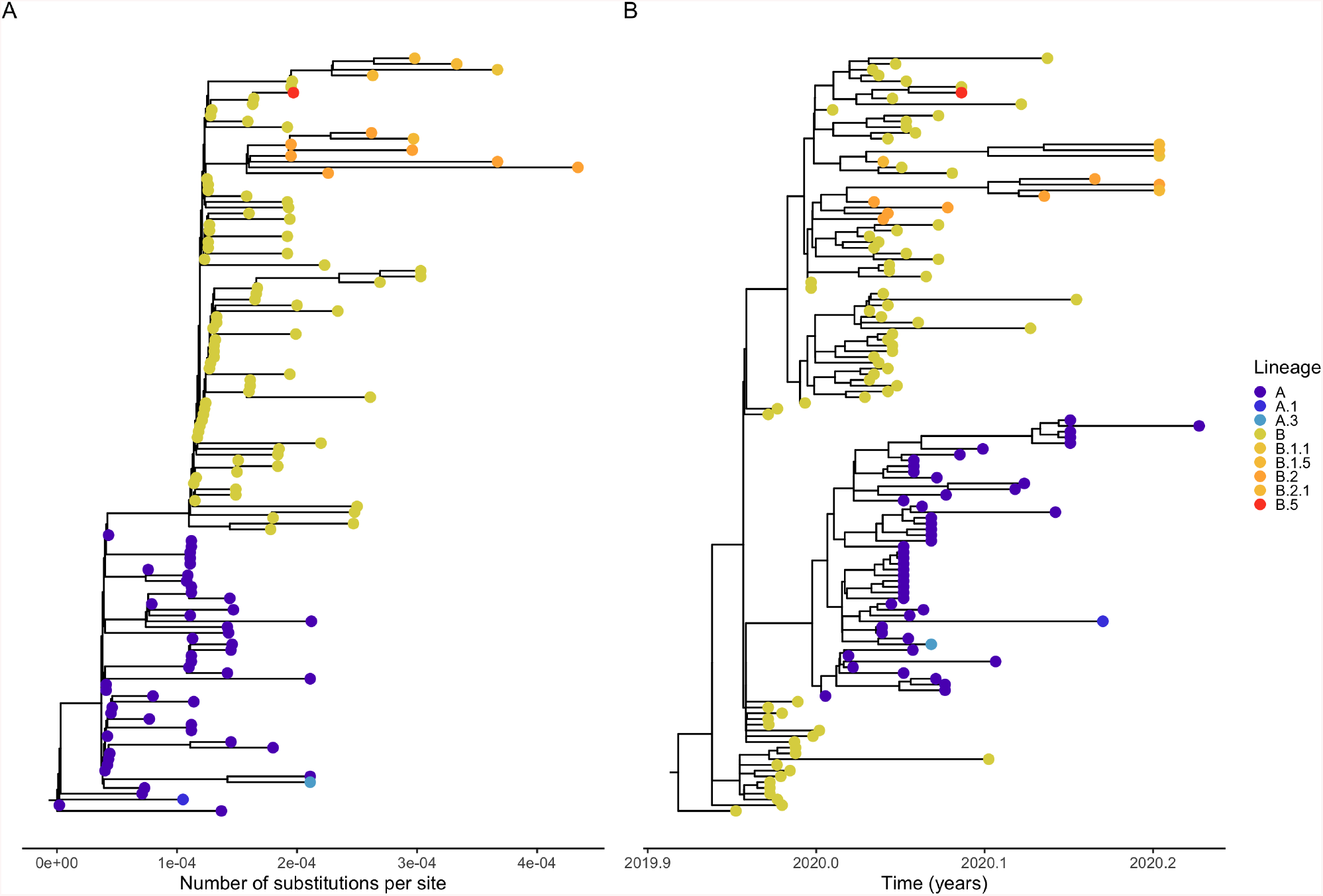
Estimated maximum likelihood (A) and estimated maximum clade credibility using tip-dating (B) of the SARS-CoV-2 phylogeny. Tips are colored according to their pangolin lineage assignment. The maximum likelihood estimate of the phylogeny was obtained using the program `RAxML-NG` (Kozlov *et al.*, 2019) under the GTR+Γ model of DNA substitution. We estimated the maximum clade credibility tree using a time-measured Bayesian phylogenetic reconstruction implemented in BEAST (Suchard *et al.*, 2018) v1.10.4. We used a GTR+Γ substitution model and the uncorrelated relaxed clock with a lognormal distribution, and specified flexible skygrid coalescent tree priors. TreeAnnotator was used to annotate the maximum clade credibility tree.

There are many different methods for inferring the root of a phylogenetic tree, but they largely depend on three possible sources of information: outgroups, the molecular clock, and non-reversibility. The latter source of information can be used if the underlying mutational process is non-reversible, that is, for some pair of nucleotides (*i*,*j*), the number of mutations from *i* to *j* differs from the number of mutations from *j* to *i*, in expectation at stationarity. However, this source of information is rarely used to root trees because it relies on strong assumptions regarding the mutational process, and it has been shown to perform poorly on real data (Huelsenbeck *et al.*, 2002). Most studies use methods based on either outgroup rooting, molecular clock rooting, or a combination of both. Outgroup rooting is perhaps the conceptually easiest method to understand, and arguably the most commonly used method. In outgroup rooting, the position in which one or more outgroups connects to the ingroup tree is the root position. Outgroup rooting can be challenged by long-branch attraction if distant outgroups are being used (e.g. Felsenstein, 1978; Graham *et al.*, 2002; Hendy and Penny, 1989; Maddison *et al.*, 1984). In such cases, the outgroup will have a tendency to be placed on the longest branches of the ingroup tree. In viruses, in particular, because of their high mutation rate, it can be challenging to identify an outgroup sequence that is sufficiently closely related to the ingroup sequences to allow reliable rooting. An alternative to outgroup rooting is molecular clock rooting, which is based on the assumption that mutations occur at an approximately constant rate, or at a rate that can be modeled and predicted using statistical models (e.g., using a relaxed molecular clock such as Drummond *et al.* (2006); Yoder and Yang (2000)). The rooting is then preferred that makes the data most compatible with the clock assumption by some criterion. Early methods for rooting using molecular clocks were often labeled midpoint rooting as some original methods were based on placing the root halfway between the most distant leaf nodes in the tree (e.g. Swofford *et al.*, 1996). More modern methods use more of the phylogenetic information, for example, by finding the rooting that minimizes the variance among leaf nodes in their distance to the root (e.g. Mai *et al.*, 2017) or produces the best linear regression of root-to-tip distances against sampling times when analyzing heterochronous data (Rambaut *et al.*, 2016). Methods for inferring phylogenetic trees that assume an ultrametric tree (i.e. a tree that perfectly follows a molecular clock), such as unweighted pair group method with arithmetic mean (UPGMA; Sokal and Michener, 1958), directly infers a rooted tree. Similarly, Bayesian phylogenetic methods using birth-death process priors (Kendall, 1948; Thompson, 1975) or coalescent priors (Kingman, 1982a, b, c) also implicitly infers the root. But even with uninformative priors on the tree the placement of the root can be estimated in Bayesian phylogenetics using molecular clock assumptions. An advantage of such methods, over methods that first infer the branch lengths of the tree and then identify the root most compatible with a molecular clock, is that they explicitly incorporate uncertainty in the branch length estimation when identifying the root and they simultaneously provide measures of statistical uncertainty in the rooting of the tree. Huelsenbeck *et al.* (2002) investigated the use of Bayesian inference of root placement and found high consistency between outgroup rooting and molecular clock rooting. The objective of this study is to determine how well the root of the SARS-CoV-2 phylogeny can be identified and to provide measures of statistical uncertainty for the placement of the root of the SARS-CoV-2 pandemic. There are several challenges when doing so. First, and most importantly, there is very little variability among the early emerging strains of the virus, challenging both molecular clock and outgroup rooting. Secondly, while the nearest outgroup sequence (RmYN02) is 97.2% identical to SARS-CoV-2 (Zhou *et al.*, 2020a), the synonymous divergence is *>*11% revealing the presence of appreciable homoplasy, providing potential additional uncertainty for outgroup rooting. Thirdly, it is unclear if a molecular clock assumption is suitable during the early phases after zoonotic transfer where selection could possibly be quite strong. Finally, coronaviruses experience substantial recombination (e.g. Boni *et al.*, 2020; Patino-Galindo *et al.*, 2020), and while there likely has not been any substantial recombination into SARS-CoV-2 since its divergence with RaTG13 and RmYN02, both of these viruses show evidence of recombination with other viruses, particularly around the gene encoding the Spike protein, that elevates the divergence from outgroup viral strains locally (e.g. Boni *et al.*, 2020; Wang *et al.*, 2020). Recombination in the outgroups is at odds with the assumption of a single phylogenetic tree shared by all sites assumed by phylogenetic models when using outgroup rooting, particularly if more than one outgroup is included in the analysis.

To investigate the possible rootings of the SARS-CoV-2 phylogeny we used six different methods and quantified the uncertainty in the placement of the root for each method on the inferred maximum likelihood topology. We note that the question of placement of a root, is a question idiosyncratic to a specific phylogeny, and to define this question we fixed the tree topology, with the exception of the root placement, in all analyses. In all cases, we applied the method to the alignment of 132 SARS-CoV-2 sequences and two putative outgroup sequences, RaTG13 and RmYN02, (see Table S1) that was constrained such that the protein-coding portions of the SARS-CoV-2 genome were in frame, and is described in detail in Wang *et al.* (2020). To ensure that we could accurately capture the rooting from available sequences, the sequences used for the analysis are chosen to be representative of the basal branches of the phylogeny and/or were early sequenced strains. There are two orders of magnitude more strains available in public databases, however these sequences are more terminally located and would provide little additional information about the placement of the root but have the potential to add a significant amount of additional noise. We are therefore focusing our efforts on the limited data set of early sequences. However, we note that future inclusion of more sequences with a basal position in the phylogeny (with few splits between the edge leading to the sequence and the root) could add additional information. The maximum likelihood estimate of the phylogeny was obtained using the program RAxML-NG (Kozlov *et al.*, 2019) under the GTR+Γ model of DNA substitution. The topology of the tree is shown in Figure 2 The outgroup sequences were pruned from the tree using nw prune from Newick utilities v1.6 (Junier and Zdobnov, 2010). Bootstrapping was preformed using the `RAxML-NG --bootstrap` option. For the RaTG13+RmYN02 analysis, only bootstrapped trees that formed a monophyletic group for RaTG13 and RmYN02 were kept. The clades of the tree were assigned according to nomenclature proposed by Rambaut *et al.* (2020) where the A and B clades are defined by the mutations 8782 and 28144 and based on whether or not they share those sites with RaTG13. The six different methods for identifying the root of the SARS-CoV-2 phylogeny were:

**FIG. 2.:**
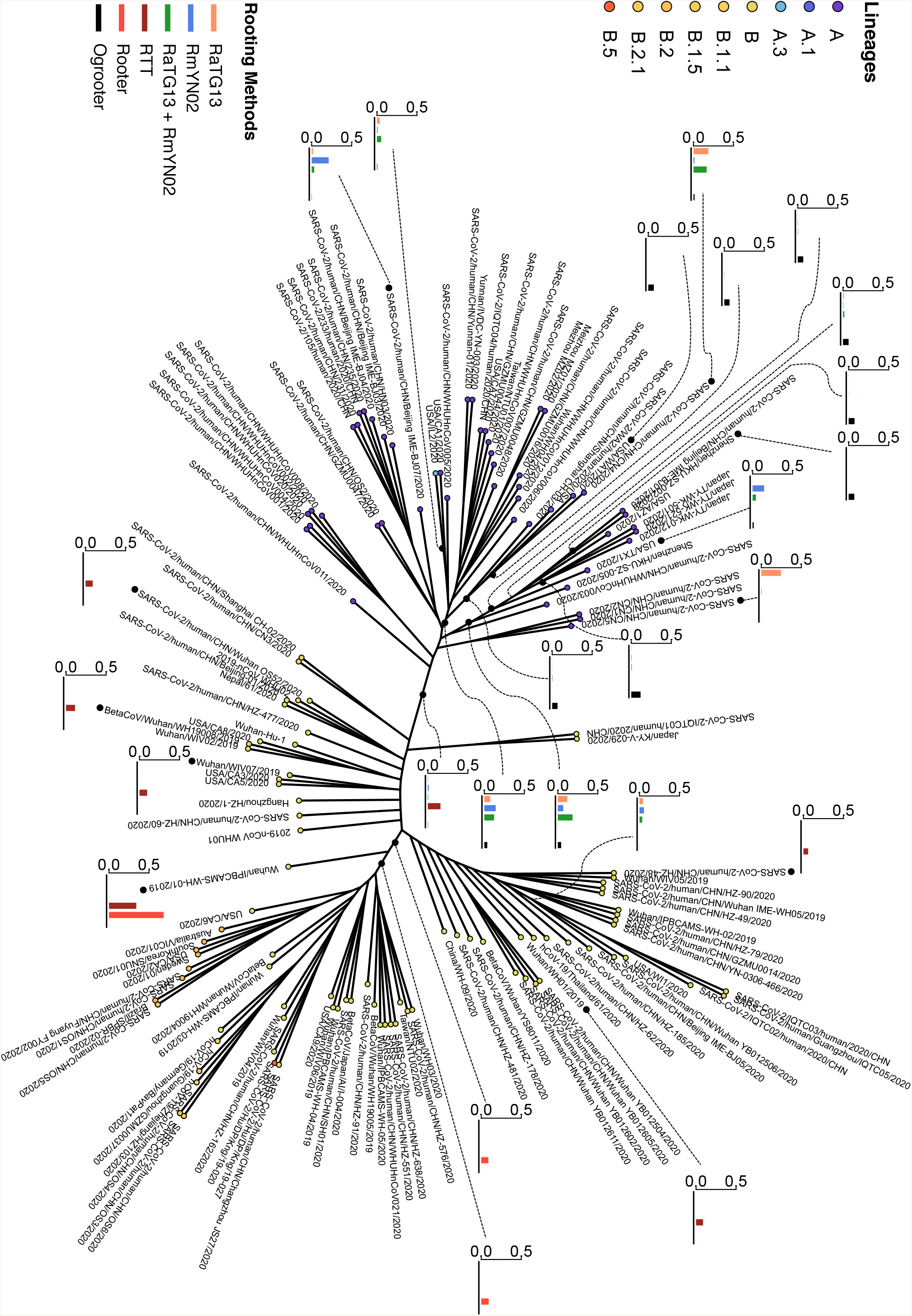
The unrooted maximum likelihood topology for the SARS-CoV-2 phylogeny of 132 genomes with probabilities for six rooting methods. Only branches with probability greater than 0.05 for at least one method are shown. Branch lengths are not to scale. The software package pangolin (https://github.com/hCoV-2019/pangolin, updated on 2020-05-01) was used for lineage assignment based on lineages updated on 2020-04-27. After running the software for assignment, only lineages called for sequences where both `aLRT` and `UFbootstrap` values which quantify the branch support in phylogeny construction are >80.

1. Outgroup rooting using RaTG13. We constrained the tree topology to be equal to the unrooted SARS-CoV-2 phylogeny, i.e. the only topological parameter estimated was the placement of the RaTG13 sequence on the unrooted SARS-CoV-2 phylogeny. We masked the potential recombination segment (NC 045512v2 positions 22851 to 23094) in RaTG13 identified in Wang *et al.* (2020) from the alignment. To quantify uncertainty we obtained 1,000 bootstrap samples. We note that while interpretation of bootstrap proportions in phylogenetics can be problematic (see Efron *et al.*, 1996), in the current context they should have a more simple interpretation as providing a confidence set for the placement of the root, i.e. if the sum of bootstrap proportions exceed 0.95 for a set of edges, under repeated sampling we would expect the root to be placed on one of these edges with probability >0.95. However, as there are very few informative sites, the bootstrap could potentially lead to poorly calibrated confidence intervals. To assess this issue, we performed 1,000 parametric simulations using pyvolve (Spielman and Wilke, 2015) using maximum likelihood estimates, from the original data set, of the model of molecular evolution and the phylogenetic tree, including branch lengths (see Table S2). For each simulation, we then estimated the tree using the same procedure as used for the real data for both the simulated data, and for 100 bootstrap replicates. We then constructed confidence sets by finding a set of branches *b* = *{b*_1_,*b*_2_,…,*b*_*k*_} such that *p*(*b*_1_)+*p*(*b*_2_)+…+*p*(*b*_*k*_) ≥ 0.95 and for which |*b*| is minimal, where *p*(*b*_*i*_) is the bootstrap proportion for branch *i*. Notice in Figure S1, that the bootstrap proportions provide rather poor measures of confidence if interpreted as such. This is likely because of the small number of mutations observed on each branch. We, therefore, consider the posterior probabilities (described in the next sections) to be more interpretable measures of uncertainty than the bootstrap proportions.
2. Outgroup rooting using RmYN02. We used the same methods as in (1) but with RmYN02 replacing RaTG13. The two potential recombination segments in RmYN02 identified in (Wang *et al.*, 2020) from the alignment (NC 045512v2 positions 21225-24252 and positions 25965-27859) were masked.
3. Outgroup rooting using both RmYN02 and RaTG13. In this case we masked all of the recombination segments identified in either RmYN02 and RaTG13 and additionally constrained the topology to make RmYN02 and RaTG13 form a clade in the unrooted phylogeny.
4. We use the ‘`rtt`’ function implemented in the R package APE (Paradis and Schliep, 2018) based on the regression method of Rambaut (2000, 2009) applied to the maximum likelihood tree. This method uses the molecular clock to root the tree. We again quantified uncertainty using 1,000 bootstrap samples.
5. We used the Bayesian molecular clock rooting method described in Huelsenbeck *et al.* (2002) but constrained to maintain the maximum likelihood topology as in the previous rooting methods. We wrote specialized software to calculate the posterior probability distribution of the root position under the molecular clock (the “Rooter” method). The program constrained the unrooted tree of the human SARS-CoV-2 sequences, estimated via maximum likelihood. However, all other parameters of the phylogenetic model were treated as random variables. The GTR+Γ model of DNA substitution was assumed in all Bayesian analyses. We used Markov chain Monte Carlo (MCMC) with 10,000,000 cycles with a sample frequency of 1,000 to update all of the model parameters. For the outgroup criterion, we initialized the tip dates using the sample dates of the viruses (which ranged from December 23, 2019 to March 24, 2020). The molecular clock was enforced, with an exponential prior with parameter *λ* = 1000 placed on the tree height.
6. We used an outgroup rooting method (the “Ogrooter” method) as described in (5) except where each branch length had an independent exponential prior with parameter *λ* = 1000. The outgroup criterion was used to root the tree. That is, we kept track of where the RaTG13 and RmYN02 sequences, which were forced to be monophyletic, joined the ingroup tree of 132 human SARS-CoV-2 sequences. We report the marginal posterior probability of the root position, which is approximated using MCMC as the fraction of the time the outgroup sequences joined the various branches.

Notice, the four methods for outgroup rooting are largely compatible (Figure 2). Most of the bootstrap replicates place the root in one of two places: in a clade leading to three Japanese sequences, two sequences from the USA, two Shenzhen sequences, and one Beijing sequence (with bootstrap proportion varying between 0.068 and 0.184, and posterior probability 0.0413) and in a clade leading to two Washington sequences, one Shanghai sequence, and one Zheijiang sequence (with bootstrap proportion varying between 0.074 and 0.142, and posterior probability 0.0363). None of these rootings are very epidemiologically plausible given that the first outbreak of SARS-CoV-2 was identified in Wuhan. There are also positive bootstrap proportions on other edges of the tree. Importantly, there is not a single placement that has high bootstrap proportion. In fact, the highest bootstrap proportion on any edge of the tree for any bootstrap method is only 0.245 and when using both RmYN02 and RaTG13, no placement has a higher bootstrap proportion than 0.2. Perhaps surprisingly, the bootstrap proportions does not get more concentrated when adding both RmYN02 and RaTG13. A possible explanation for this is the reduction in alignment length when removing the recombination fragments from RmYN02. The two methods for placing the root using a molecular clock are also mostly compatible with each other. Rooter places about half of the posterior probability (0.464), and the root-to-tip regression rooting method (rtt) places 0.341 bootstrap proportion, at the earliest collected sequence from Wuhan (Wuhan/IPBCAMS-WH-01/2019). Rooter also places 0.137 probability on the edge leading to this sequence and 0.151 probability on the sister edge to this sequence. However, there is also considerable probability assigned in various other positions. No singular placement in the tree receives more than 0.464 probability.

To investigate differences in signs of temporal signal for the outgroup rooting and the molecular clock rooting, we calculated root-to-tip distances using TempEst v1.5.3 (Rambaut *et al.*, 2016) for the ML tree using the outgroup rooting (Fig S2) and a re-rooting of the ML tree using the molecular clock rooting (Fig S3). Re-rooting was performed using `nw_reroot` from Newick utilities v1.6 (Junier and Zdobnov, 2010). As expected, the molecular clock rooting has more temporal signal (r=0.403, value=7.226 ×10^−8^) than the outgroup rooting (r=0.271, p-value=3.367×10^−4^). Additionally, we infer the root age of the molecular clock rooting to be in mid-October 2019 (2019.794) with a 95% confidence interval of [2019.225, 2020.363] and we estimate the rate of evolution to be 5.470×10^−4^ substitutions per site per year with a 95% confidence interval of [3.049×10^−4^, 7.891×10^−4^]. Despite our small sample size and our focus on basal lineages of the SARS-CoV-2 tree to assess uncertainty in the rooting of the phylogeny, this is largely compatible with other estimates of the time to the most common recent ancestor (e.g. Lai *et al.*, 2020; van Dorp *et al.*, 2020). The inferred root age of the outgroup rooting is much earlier than other estimates, in mid-August of 2019 (2019.632), with a much wider 95% confidence interval of [2017.632, 2020.469], an indication of greater uncertainty and less molecular clock signal. We estimate the rate of evolution for the outgroup rooting to be 3.547 ×10^−4^ substitutions per site per year with a 95% confidence interval of [1.112×10^−4^, 5.969×10^−4^]. We calculated the confidence intervals for the root age and the rate of evolution using the standard errors of the x-intercept and the slope, respectively, which was estimated using a nonparametric bootstrap of the alignment sites with *n* = 5,000. Using the same bootstrapping of the alignment, we calculated the p-values for the correlations using a two-sided Wald test.

We also assessed the temporal signal in the data using Bayesian evaluation of temporal signal (BETS) (Duchene *et al.*, 2020a), which estimates the marginal likelihoods using a model that contains sampling times and a model containing no sampling times. A model is preferred over another according to their ratio of marginal likelihoods. The model using sampling times is expected to have the highest statistical fit if the data contains temporal signal. BETS provides further evidence that there is temporal signal in the data with only a modest improvement in statistical fit for the relaxed clock over the strict clock model (Fig S4). Additionally, Duchene *et al.* (2020b) have used BETS to show that there is positive evidence for temporal signal if genomes past February 2nd are included in the data. While molecular clock rooting and the outgroup rooting strategies internally give qualitatively similar results, they are largely incompatible with each other. The molecular clock rooting places the root in the B clade with high confidence, while outgroup rooting places the root in the A clade with similarly high confidence. The reason for this discrepancy is unclear, but it could be caused either by deviations from a molecular clock or excess back-mutations, i.e. unexpectedly many mutations in the same site occurring both within the SARS-CoV-2 phylogeny and on the lineage leading to the outgroup(s). We were able to capture outgroup rooting compatible with the molecular clock rooting (obtaining an outgroup rooting in clade B instead of clade A) by removing three positions from the alignment (8782, 18060, and 28144) (Figure S5). All of these positions have negative phyloP values based on the UCSC 119way alignment (Fernandes *et al.*, 2020), which suggests fast evolution. While positions 8782 and 18060 are synonymous changes, position 28144 is a missense mutation in orf8 whose function is unclear, but which has also back-mutated in more recent samples of the A clade. Also, all three mutations are between T and C which occur with particularly high rate within SARS-CoV-2 (see e.g., https://virological.org/t/issues-with-sars-cov-2-sequencing-data/473). The most likely explanation for the observed discrepancy between the rootings might be hypermutatibility in these sites causing excess back-mutations, suggesting that the molecular clock rooting is more reliable. However, we cannot exclude an increased rate of mutation (or sequencing errors) in the A clade that would attract the root to this clade. However, both methods of rooting reveal substantial uncertainty in the placement of the root.

The rooting of the SARS-CoV-2 phylogeny has important implications for our understanding of Covid19 epidemiology. Rooting in the B clade is compatible with an origin in or near Wuhan, while a rooting in Clade A suggests alternative origins of the virus, perhaps outside East Asia. The phylogenetic evidence alone, which is the focus of our study, is not sufficient to resolve this issue. However, we note that the vast majority of epidemiological evidence points to an origin of the virus in or near Wuhan. Furthermore, hypermutation in a few sites, not accounted for in standard models of molecular evolution, might be able to explain the signal in the outgroup rooting towards a root in Clade A. For that reason we consider the rooting in Clade B, as estimated in molecular clock analyses, and implied in standard phylodynamic analyses (e.g. Lemey *et al.*, 2020), to be the most plausible rooting. However, it might be prudent to avoid strong inferences regarding the early divergence of SARS-CoV-2 based on a fixed rooting in either the A or the B clade, and analyses based on the outgroup rooting should be avoided until outgroups more closely related to SARS-CoV-2 have been discovered. While the outgroup rooting seems the least compatible with the available epidemiological evidence, we recommend that studies that use the rooting for inferences: (1) use methods that can take uncertainty in the rooting into account, for example by integrating over possible rootings, as accomplished by Bayesian phylogenetic methods, and (2) combine evidence from outgroup rooting and molecular clock rootings. The latter could, for example, be accomplished in a Bayesian framework by also including outgroup sequences in traditional phylodynamic analyses, but with a different prior governing the evolution and epidemiology of the outgroup sequences than that used for the ingroup sequences.

## Online Resources

Rooter and Ogrooter are available for download at https://github.com/NielsenBerkeleyLab/rooter and https://github.com/NielsenBerkeleyLab/ogrooter, respectively.

## Acknowledgments

We thank Dr. Adi Stern for discussion. The research was funded by Koret-UC Berkeley-Tel Aviv University Initiative in Computational Biology and Bioinformatics to RN.

## References

Benvenuto, D., Giovanetti, M., Salemi, M., Prosperi, M., De Flora, C., Junior Alcantara, L. C., Angeletti, S., and Ciccozzi, M. 2020. The global spread of 2019-ncov: a molecular evolutionary analysis. Pathogens and Global Health, 114(2): 64–67.

Boni, M. F., Lemey, P., Jiang, X., Lam, T. T.-Y., Perry, B., Castoe, T., Rambaut, A., and Robertson, D. L. 2020. Evolutionary origins of the sars-cov-2 sarbecovirus lineage responsible for the covid-19 pandemic. bioRxiv.

Drummond, A. J., Ho, S. Y., Phillips, M. J., and Rambaut, 2006. Relaxed phylogenetics and dating with confidence. PLoS biology, 4(5).

Duchene, S., Lemey, P., Stadler, T., Ho, S. Y. W., Duchene, D. A., Dhanasekaran, V., and Baele, G. 2020a. Bayesian Evaluation of Temporal Signal in Measurably Evolving Populations. Molecular Biology and Evolution. msaa163.

Duchene, S., Featherstone, L., Haritopoulou-Sinanidou, M., Rambaut, A., Lemey, P., and Baele, G. 2020b. Temporal signal and the phylodynamic threshold of sars-cov-2. bioRxiv.

Efron, B., Halloran, E., and Holmes, S. 1996. Bootstrap confidence levels for phylogenetic trees. Proceedings of the National Academy of Sciences, 93(23): 13429–13429.

Felsenstein, J. 1978. Cases in which parsimony or compatibility methods will be positively misleading. Systematic zoology, 27(4): 401–410.

Fernandes, J. D., Hinrichs, A. S., Clawson, H., Navarro Gonzalez, J., Lee, B. T., Nassar, L. R., Raney, B. J., Rosenbloom, K. R., Nerli, S., Rao, A., Schmelter, D., Zweig, A. S., Lowe, T. M., Ares, M., Corbet-Detig, R., Kent, W. J., Haussler, D., and Haeussler, M. 2020. The ucsc sars-cov-2 genome browser. bioRxiv.

Giovanetti, M., Benvenuto, D., Angeletti, S., and Ciccozzi, M. 2020. The first two cases of 2019-ncov in italy: Where they come from? Journal of medical virology, 92(5): 518–521.

Gomez-Carballa, A., Bello, X., Pardo-Seco, J., Martinon-Torres, F., and Salas, A. 2020. Mapping genome variation of sars-cov-2 worldwide highlights the impact of covid-19 super-spreaders. Genome Research, pages gr–266221.

Graham, S. W., Olmstead, R. G., and Barrett, S. C. 2002. Rooting phylogenetic trees with distant outgroups: a case study from the commelinoid monocots. Molecular biology and evolution, 19(10): 1769–1781.

Hendy, M. D. and Penny, D. 1989. A framework for the quantitative study of evolutionary trees. Systematic zoology, 38(4): 297–309.

Huelsenbeck, J. P., Bollback, J. P., and Levine, A. M. 2002. Inferring the root of a phylogenetic tree. Systematic biology, 51(1): 32–43.

Junier, T. and Zdobnov, E. M. 2010. The newick utilities: high-throughput phylogenetic tree processing in the unix shell. Bioinformatics, 26(13): 1669–1670.

Kendall, D. G. 1948. On the generalized “birth-death” process. Annual Review of Mathematical Statistics, 19: 1–15.

Kingman, J. F. C. 1982a. The coalescent. Stochastic Processes and their Applications, 13: 235–248.

Kingman, J. F. C. 1982b. Exchangeability and the evolution of large populations. In G. Koch and F. Spizzichino, editors, Exchangeability in Probability and Statistics, pages 97–112. North-Holland.

Kingman, J. F. C. 1982c. On the genealogy of large populations. In J. Gani and E. J. Hannan, editors, Essays in Statistical Science: Papers in Honour of P. A. P. Moran, Journal of Applied Probability, Special Volume 19A, pages 27–43. Applied Probability Trust.

Kozlov, A. M., Darriba, D., Flouri, T., Morel, B., and Stamatakis, A. 2019. Raxml-ng: a fast, scalable and user-friendly tool for maximum likelihood phylogenetic inference. Bioinformatics, 35(21): 4453–4455.

Lai, A., Bergna, A., Acciarri, C., Galli, M., and Zehender, G. 2020. Early phylogenetic estimate of the effective reproduction number of sars-cov-2. Journal of medical virology, 92(6): 675–679.

Lemey, P., Hong, S., Hill, V., Baele, G., Poletto, C., Colizza, V., O’Toole, A., McCrone, J. T., Andersen, K. G., Worobey, M., et al. 2020. Accommodating individual travel history, global mobility, and unsampled diversity in phylogeography: a sars-cov-2 case study. bioRxiv.

Li, J., Li, Z., Cui, X., and Wu, C. 2020a. Bayesian phylodynamic inference on the temporal evolution and global transmission of sars-cov-2. The Journal of infection.

Li, Q. et al. 2020b. An outbreak of ncip (2019-ncov) infection in chinawuhan, hubei province, 2019-2020. China CDC Weekly, 2(5): 79–80.

Li, X., Zai, J., Zhao, Q., Nie, Q., Li, Y., Foley, B. T., and Chaillon, A. 2020c. Evolutionary history, potential intermediate animal host, and cross-species analyses of sars-cov-2. Journal of medical virology, 92(6): 602–611.

Li, X., Zai, J., Wang, X., and Li, Y. 2020d. Potential of large first generation human-to-human transmission of 2019-ncov. Journal of medical virology, 92(4): 448–454.

Maddison, W. P., Donoghue, M. J., and Maddison, D. R. 1984. Outgroup analysis and parsimony. Systematic biology, 33(1): 83–103.

Mai, U., Sayyari, E., and Mirarab, S. 2017. Minimum variance rooting of phylogenetic trees and implications for species tree reconstruction. PloS one, 12(8).

Morel, B., Barbera, P., Czech, L., Bettisworth, B., Huebner, L., Lutteropp, S., Serdari, D., Kostaki, E.-G., Mamais, I., Kozlov, A., et al. 2020. Phylogenetic analysis of sars-cov-2 data is difficult. bioRxiv.

Nie, Q., Li, X., Chen, W., Liu, D., Chen, Y., Li, H., Li, D., Tian, M., Tan, W., and Zai, J. 2020. Phylogenetic and phylodynamic analyses of sars-cov-2. Virus research, 287: 198098.

Paradis, E. and Schliep, K. 2018. ape 5.0: an environment for modern phylogenetics and evolutionary analyses in R. Bioinformatics, 35(3): 526–528.

Patino-Galindo, J. A., Filip, I., AlQuraishi, M., and Rabadan, R. 2020. Recombination and lineage-specific mutations led to the emergence of sars-cov-2. bioRxiv.

Rambaut, A. 2000. Estimating the rate of molecular evolution: incorporating non-contemporaneous sequences into maximum likelihood phylogenies. Bioinformatics, 16(4): 395–399.

Rambaut, A. 2009. Path-o-gen: temporal signal investigation tool.

Rambaut, A., Lam, T. T., Max Carvalho, L., and Pybus, O. G. 2016. Exploring the temporal structure of heterochronous sequences using tempest (formerly path-o-gen). Virus evolution, 2(1): vew007.

Rambaut, A., Holmes, E. C., Hill, V., OToole, A., McCrone, J., Ruis, C., du Plessis, L., and Pybus, O. 2020. A dynamic nomenclature proposal for sars-cov-2 to assist genomic epidemiology. bioRxiv.

Shen, L., Bard, J. D., Biegel, J. A., Judkins, A. R., and Gai, X. 2020. Comprehensive genome analysis of 6,000 usa sars-cov-2 isolates reveals haplotype signatures and localized transmission patterns by state and by country. medRxiv.

Sokal, R. R. and Michener, C. D. 1958. A statistical method for evaluating systematic relationships. Univ. Kans. Sci. Bull., 28: 1409–1438.

Spielman, S. J. and Wilke, C. O. 2015. Pyvolve: a flexible python module for simulating sequences along hylogenies. PloS one, 10(9): e0139047.

Suchard, M. A., Lemey, P., Baele, G., Ayres, D. L., Drummond, A. J., and Rambaut, A. 2018. Bayesian phylogenetic and phylodynamic data integration using beast 1.10. Virus evolution, 4(1): vey016.

Swofford, D., Olsen, P., Waddell, P., and Hillis, D. 1996. Molecular systematics, chapter phylogenetic inference. Sinauer Associates, 15: 407–514.

Tang, X., Wu, C., Li, X., Song, Y., Yao, X., Wu, X., Duan, Y., Zhang, H., Wang, Y., Qian, Z., Cui, J., and Lu, J. 2020. On the origin and continuing evolution of SARS-CoV-2. National Science Review. nwaa036.

Thompson, E. A. 1975. Human Evolutionary Trees. Cambridge University Press, Cambridge, England.

van Dorp, L., Acman, M., Richard, D., Shaw, L. P., Ford, C. E., Ormond, L., Owen, C. J., Pang, J., Tan, C. C., Boshier, F. A., et al. 2020. Emergence of genomic diversity and recurrent mutations in sars-cov-2. Infection, Genetics and Evolution, page 104351.

Wang, H., Pipes, L., and Nielsen, R. 2020. Synonymous mutations and the molecular evolution of sars-cov-2 origins. bioRxiv.

World Health Organization 2020. Coronavirus disease 2019 (covid-19) situation report 41. https://www.who.int/docs/default-source/coronaviruse/situation-reports/20200301-sitrep-41-covid-19.pdf?sfvrsn=6768306d_2.

Wu, F., Zhao, S., Yu, B., Chen, Y.-M., Wang, W., Song, Z.-G., Hu, Y., Tao, Z.-W., Tian, J.-H., Pei, Y.-Y., et al. 2020. A new coronavirus associated with human respiratory disease in china. Nature, 579(7798): 265–269.

Yoder, A. D. and Yang, Z. 2000. Estimation of primate speciation dates using local molecular clocks. Molecular Biology and Evolution, 17(7): 1081–1090.

Yu, W.-B., Tang, G.-D., Zhang, L., and Corlett, R. T. 2020. Decoding the evolution and transmissions of the novel pneumonia coronavirus (sars-cov-2/hcov-19) using whole genomic data. Zoological Research, 41(3): 247.

Zhang, L., Shen, F.-m., Chen, F., and Lin, Z. 2020. Origin and evolution of the 2019 novel coronavirus. Clinical Infectious Diseases.

Zhou, H., Chen, X., Hu, T., Li, J., Song, H., Liu, Y., Wang, P., Liu, D., Yang, J., Holmes, E. C., et al. 2020a. A novel bat coronavirus reveals natural insertions at the s1/s2 cleavage site of the spike protein and a possible recombinant origin of hcov-19. Current Biology.

Zhou, P., Yang, X.-L., Wang, X.-G., Hu, B., Zhang, L., Zhang, W., Si, H.-R., Zhu, Y., Li, B., Huang, C.-L., et al. 2020b. A pneumonia outbreak associated with a new coronavirus of probable bat origin. nature, 579(7798): 270–273.

