## Supplementary Material for "Assessing uncertainty in the rooting of the SARS-CoV-2 phylogeny"

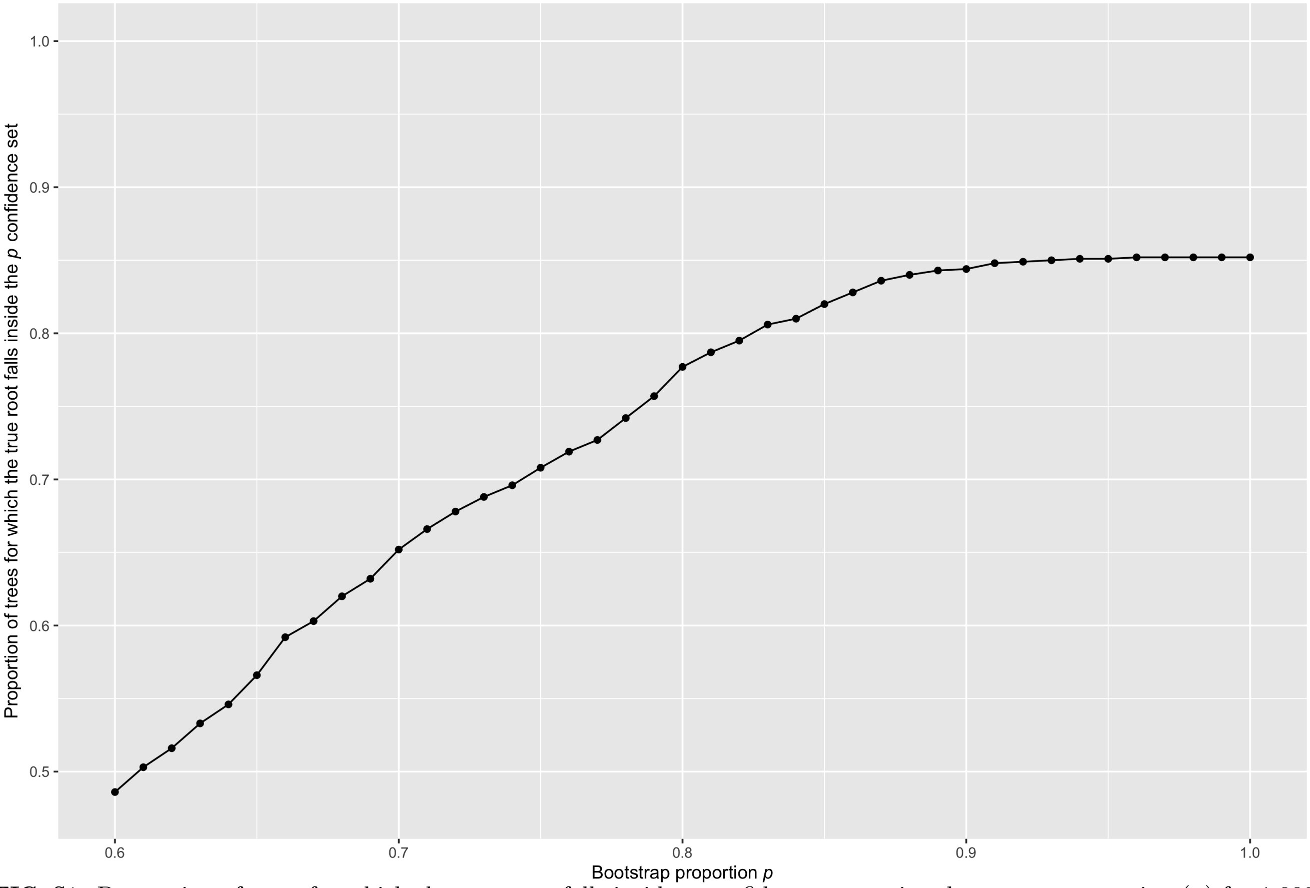

**FIG. S1.** Proportion of trees for which the true root falls inside $p$ confidence set against bootstrap proportion ($p$) for 1,000 parametric simulations. Simulations were performed with pyvolve (Spielman and Wilke, 2015) using maximum likelihood estimates, from the original data set, of the model of molecular evolution and the phylogenetic tree, including branch lengths (see Table S2).

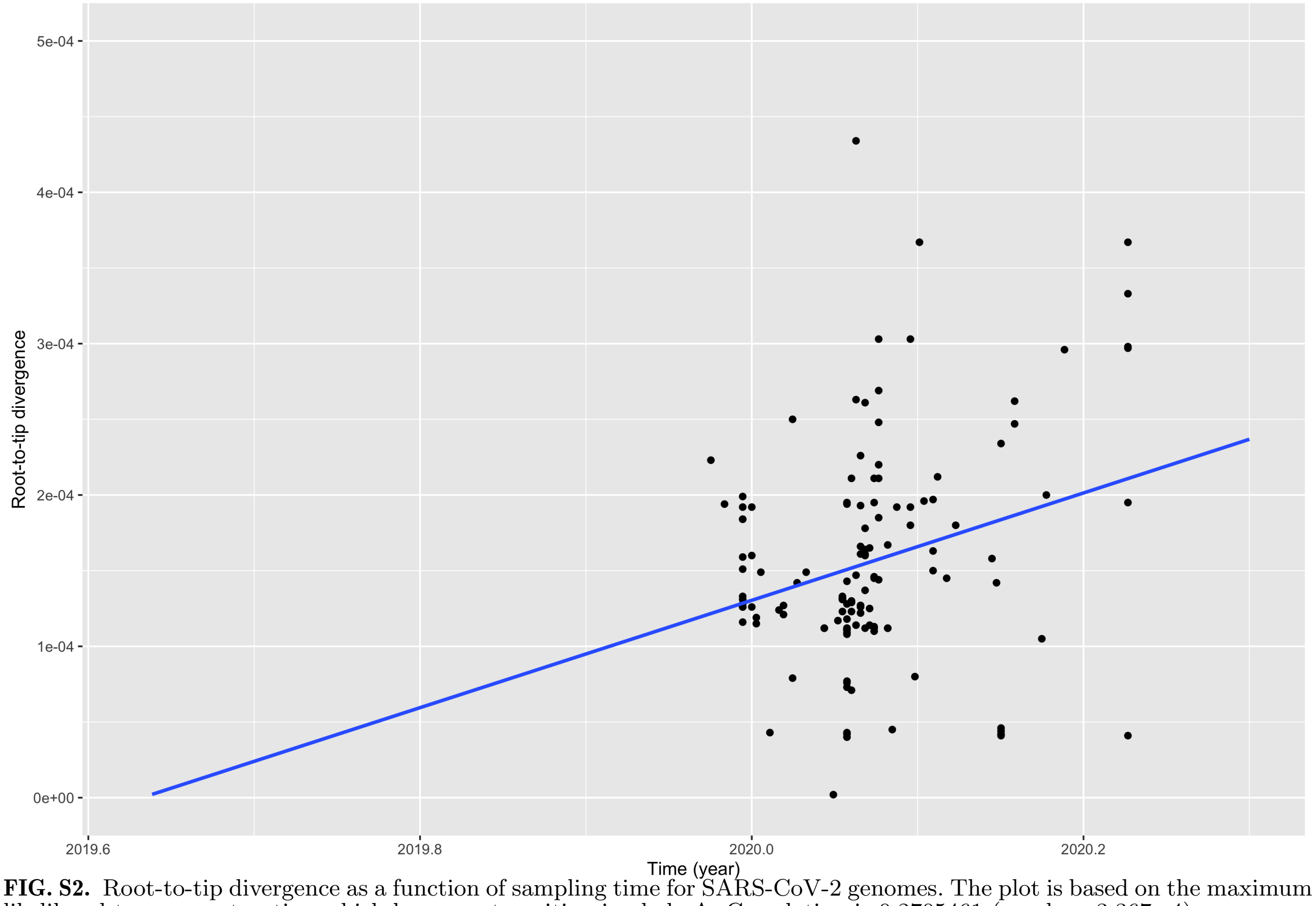


**FIG. S2.** Root-to-tip divergence as a function of sampling time for SARS-CoV-2 genomes. The plot is based on the maximum likelihood tree reconstruction which has a root position in clade A. Correlation is 0.2705461 (p-value=3.367e-4).

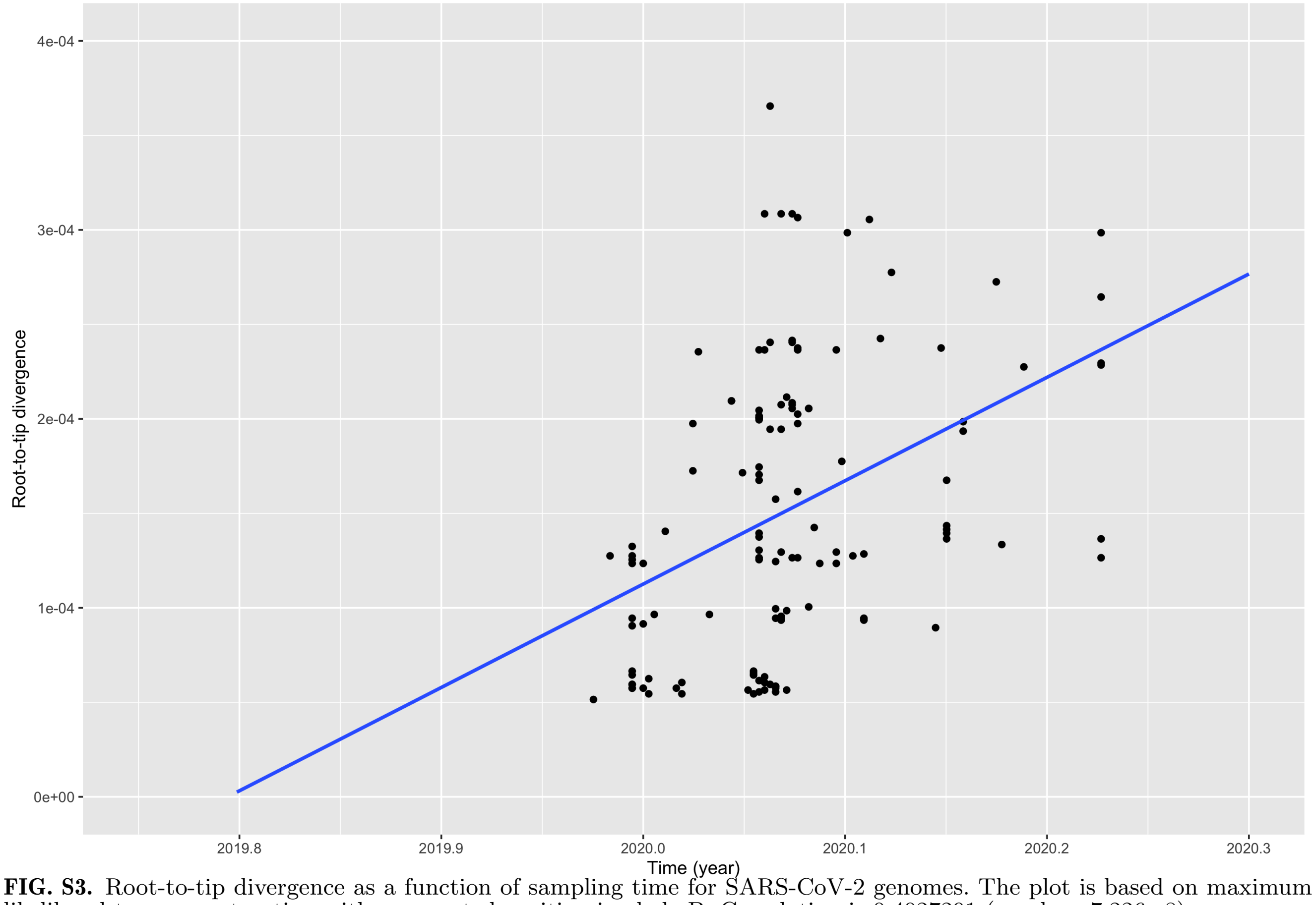


**FIG. S3.** Root-to-tip divergence as a function of sampling time for SARS-CoV-2 genomes. The plot is based on maximum likelihood tree reconstruction with a re-rooted position in clade B. Correlation is 0.4027291 (p-value=7.226e-8).

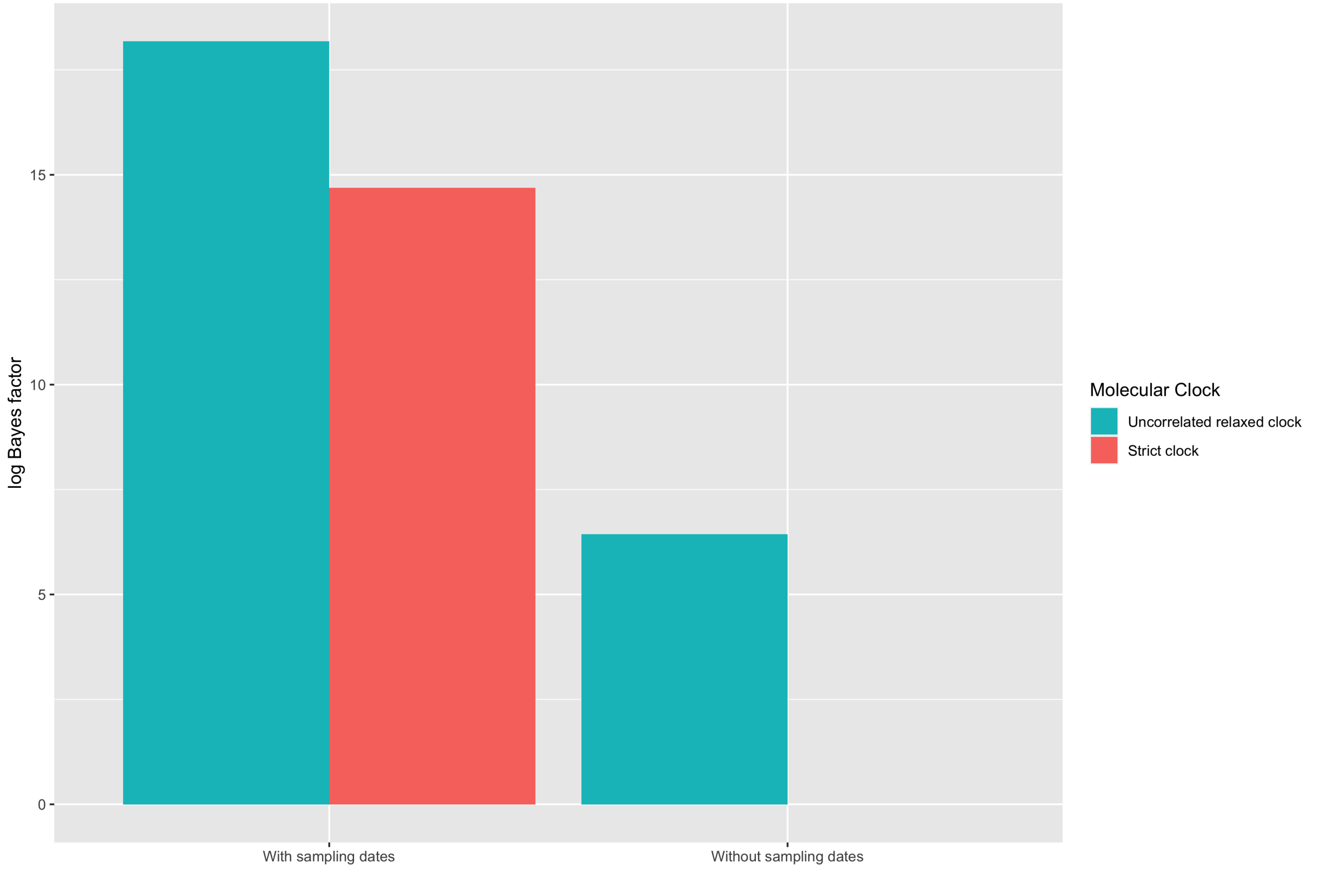


**FIG. S4.** Bayesian evaluation of temporal signal (BETS) results where the poorest performing model has a log Bayes factor of 0. We analyzed the sequences in Table S1 using BEAST 1.10.4 using a Markov chain Monte Carlo (MCMC) length of 10000000 steps and sampling every 1000 steps. We compared the statistical fit of two molecular clock models, strict clock and uncorrelated lognormal relaxed clock, with and without sampling times. For each combination of molecular clock and sampling times, we calculated the (log) marginal likelihood using generalized stepping stone sampling (Baele *et al.*, 2016), for which we used 200 path steps with a chain length for each 100000 iterations.

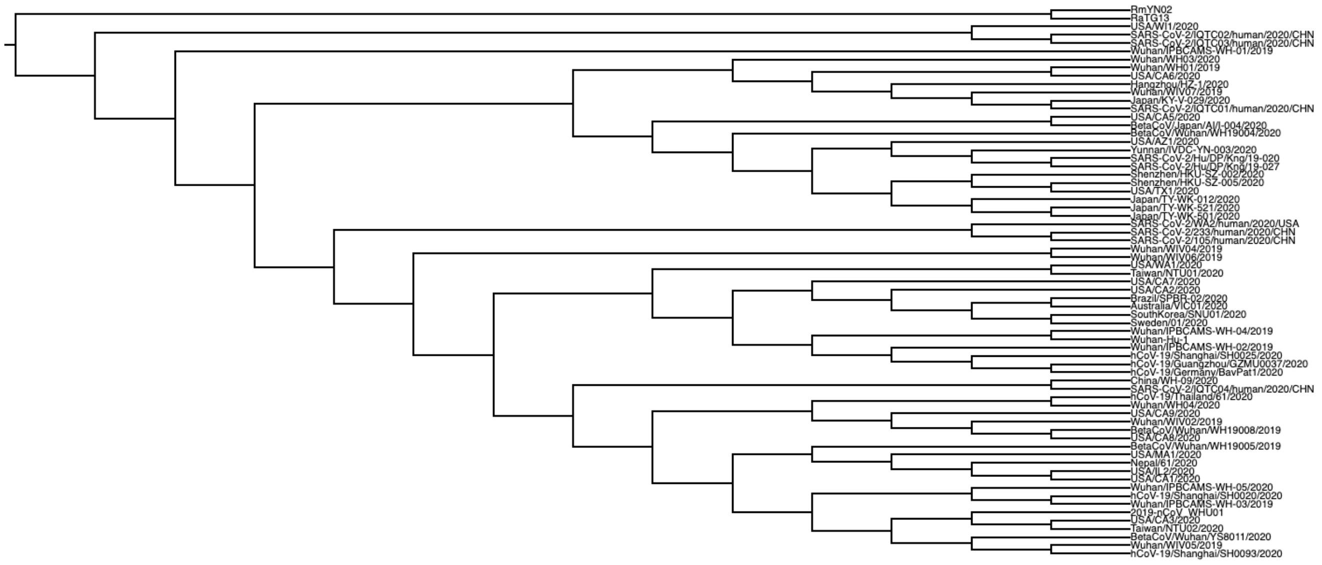


**FIG. S5.** Outgroup rooting by removal of positions 8782, 18060, and 28144 from the alignment.

Table S1.: Accessions and sources of the SARS-CoV-2 genome sequences used in this study.

| Genome Name | Source | Accession ID |
|---|---|---|
| Wuhan/IPBCAMS-WH-01/2019 | GenBank | MT019529 |
| BetaCoV/Wuhan/WH-01/2019 | GenBank | LR757998 |
| Wuhan/WIV02/2019 | GenBank | MN996527 |
| Wuhan/WIV04/2019 | GenBank | MN996528 |
| Wuhan/WIV05/2019 | GenBank | MN996529 |
| Wuhan/WIV06/2019 | GenBank | MN996530 |
| Wuhan/WIV07/2019 | GenBank | MN996531 |
| Wuhan/IPBCAMS-WH-02/2019 | GenBank | MT019530 |
| Wuhan/IPBCAMS-WH-03/2019 | GenBank | MT019531 |
| Wuhan/IPBCAMS-WH-04/2019 | GenBank | MT019532 |
| BetaCoV/Wuhan/WH19008/2019 | NMDC | NMDC60013002_06 |
| BetaCoV/Wuhan/WH19005/2019 | NMDC | NMDC60013002_10 |
| 2019-nCoV_HKU-SZ-002a_2020 | GenBank | MN938384 |
| 2019-nCoV_HKU-SZ-005b_2020 | GenBank | MN975262 |
| SouthKorea/SNU01/2020 | GenBank | MT039890 |
| BetaCoV/Wuhan/WH-03/2019 | GenBank | LR757996 |
| Wuhan/IPBCAMS-WH-05/2020 | GenBank | MT019533 |
| BetaCoV/Wuhan/WH19004/2020 | NMDC | NMDC60013002_09 |
| 2019-nCoV_WHU01 | GenBank | MN988668 |
| Wuhan/WH04/2019 | GenBank | LR757995 |
| BetaCoV/Wuhan/YS8011/2020 | NMDC | NMDC60013002_07 |
| China/WH-09/2020 | GenBank | MT093631 |
| Nepal/61/2020 | GenBank | MT072688 |
| SARS-CoV-2/human/CHN/Yunnan-01/2020 | GenBank | MT049951 |
| USA/WA1/2020 | GenBank | MN985325 |
| Hangzhou/HZ-1/2020 | GenBank | MT039873 |
| USA/CA2/2020 | GenBank | MN994468 |

| | | |
|---|---|---|
| USA-AZ1/2020 | GenBank | MN997409 |
| USA/CA1/2020 | GenBank | MN994467 |
| BetaCoV/Japan/AI/I-004/2020 | GenBank | LC521925 |
| Australia/VIC01/2020 | GenBank | MT007544 |
| SARS-CoV-2/105/human/2020/CHN | GenBank | MT135041 |
| USA/CA6/2020 | GenBank | MT044258 |
| USA/IL2/2020 | GenBank | MT044257 |
| SARS-CoV-2/233/human/2020/CHN | GenBank | MT135043 |
| Japan/KY/V-029/2020 | GenBank | LC522972 |
| Japan/TY/WK-012/2020 | GenBank | LC522973 |
| USA/CA3/2020 | GenBank | MT027062 |
| USA/CA5/2020 | GenBank | MT027064 |
| SARS-CoV-2/IQTC02/human/2020/CHN | GenBank | MT123291 |
| Japan/TY/WK-501/2020 | GenBank | LC522974 |
| Japan/TY/WK-521/2020 | GenBank | LC522975 |
| USA/WI1/2020 | GenBank | MT039887 |
| Taiwan/NTU01/2020 | GenBank | MT066175 |
| Taiwan/NTU02/2020 | GenBank | MT066176 |
| SARS-CoV-2/IQTC01/human/2020/CHN | GenBank | MT123290 |
| USA/CA7/2020 | GenBank | MT106052 |
| Sweden/01/2020 | GenBank | MT093571 |
| SARS-CoV-2/Hu/DP/Kng/19-020 | GenBank | LC528232 |
| SARS-CoV-2/Hu/DP/Kng/19-027 | GenBank | LC528233 |
| USA/CA8/2020 | GenBank | MT106053 |
| USA/CA9/2020 | GenBank | MT118835 |
| SARS-CoV-2/IQTC04/human/2020/CHN | GenBank | MT123292 |
| SARS-CoV-2/IQTC03/human/2020/CHN | GenBank | MT123293 |
| Brazil/SPBR-02/2020 | GenBank | MT126808 |
| SARS-CoV-2/WA2/human/2020/USA | GenBank | MT152824 |
| Wuhan-Hu-1 | GenBank | NC_045512 |

| | | |
|---|---|---|
| hCoV-19/Thailand/61/2020 | GISAID | EPI_ISL_403962 |
| hCoV-19/Germany/BavPat1/2020 | GISAID | EPI_ISL_406862 |
| hCoV-19/Guangzhou/GZMU0037/2020 | GISAID | EPI_ISL_416334 |
| hCoV-19/Shanghai/SH0025/2020 | GISAID | EPI_ISL_416334 |
| hCoV-19/Zhejiang/HZ103/2020 | GISAID | EPI_ISL_422425 |
| SARS-CoV-2/human/CHN/Beijing_IME-BJ01/2020 | GenBank | MT291831.1 |
| USA/TX1/2020 | GenBank | MT106054 |
| SARS-CoV-2/human/CHN/WHUHnCoV003/2020 | GenBank | MT079845 |
| SARS-CoV-2/human/CHN/CN2/2020 | GenBank | MT407650 |
| SARS-CoV-2/human/CHN/CN1/2020 | GenBank | MT407649 |
| SARS-CoV-2/human/CHN/CN5/2020 | GenBank | MT407651 |
| SARS-CoV-2/IQTC01/human/2020/CHN | GenBank | MT123290 |
| SARS-CoV-2/human/CHN/HZ-477/2020 | GenBank | MT253699 |
| SARS-CoV-2/human/CHN/HZ-178/2020 | GenBank | MT253697 |
| SARS-CoV-2/human/CHN/HZ-62/2020 | GenBank | MT253706 |
| SARS-CoV-2/human/CHN/HZ-90/2020 | GenBank | MT253709 |
| SARS-CoV-2/human/CHN/HZ-48/2020 | GenBank | MT253701 |
| SARS-CoV-2/human/CHN/Wuhan_IME-WH05/2019 | GenBank | MT291830 |
| SARS-CoV-2/human/CHN/HZ-49/2020 | GenBank | MT253702 |
| SARS-CoV-2/human/CHN/HZ-185/2020 | GenBank | MT253698 |
| SARS-CoV-2/human/CHN/HZ-79/2020 | GenBank | MT253708 |
| SARS-CoV-2/human/CHN/GZMU0014/2020 | GenBank | MT568634 |
| SARS-CoV-2/human/CHN/YN-0306-466/2020 | GenBank | MT396241 |
| SARS-CoV-2/human/CHN/Beijing_IME-BJ05/2020 | GenBank | MT291835 |
| SARS-CoV-2/human/Guangzhou/IQTC05/2020 | GenBank | MT446312 |
| SARS-CoV-2/human/CHN/Wuhan$_{Y}B$012506/2020 | GenBank | MT259230 |
| Wuhan/WH01/2019 | GenBank | MT291826 |
| SARS-CoV-2/human/CHN/Wuhan_YB012602/2020 | GenBank | MT259229 |
| SARS-CoV-2/human/CHN/Wuhan_YB012504/2020 | GenBank | MT259231 |
| SARS-CoV-2/human/CHN/Wuhan_YB012605/2020 | GenBank | MT259228 |

| | | |
|---|---|---|
| SARS-CoV-2/human/CHN/Wuhan_YB012611/2020 | GenBank | MT259227 |
| SARS-CoV-2/human/CHN/HZ-481/2020 | GenBank | MT253700 |
| SARS-CoV-2/human/CHN/HZ-91/2020 | GenBank | MT253710 |
| Wuhan/WH03/2020 | GenBank | MT291828 |
| SARS-CoV-2/human/CHN/HZ-576/2020 | GenBank | MT253704 |
| SARS-CoV-2/human/CHN/HZ-638/2020 | GenBank | MT253707 |
| SARS-CoV-2/human/CHN/HZ-551/2020 | GenBank | MT253703 |
| SARS-CoV-2/human/CHN/WHUHnCoV021/2020 | GenBank | MT079854 |
| SARS-CoV-2/human/CHN/SH01/2020 | GenBank | MT121215 |
| SARS-CoV-2/human/CHN/HZ-162/2020 | GenBank | MT253696 |
| SARS-CoV-2/human/CHN/Changzhou_JS27/2020 | GenBank | MT534630 |
| hCoV-19/Germany/BavPat1/2020 | GenBank | MT270101 |
| SARS-CoV-2/human/CHN/OS6/2020 | GenBank | MT407656 |
| SARS-CoV-2/human/CHN/OS3/2020 | GenBank | MT407657 |
| SARS-CoV-2/human/CHN/OS4/2020 | GenBank | MT407659 |
| SARS-CoV-2/human/CHN/Fuyang_FY002/2020 | GenBank | MT281577 |
| SARS-CoV-2/human/CHN/OS5/2020 | GenBank | MT407655 |
| SARS-CoV-2/human/CHN/OS1/2020 | GenBank | MT407654 |
| SARS-CoV-2/human/CHN/HZ-60/2020 | GenBank | MT253705 |
| SARS-CoV-2/human/CHN/Wuhan_OS52/2020 | GenBank | MT259226 |
| 2019-nCoV_WHU02 | GenBank | MN988669 |
| SARS-CoV-2/human/CHN/Beijing-01/2020 | GenBank | MT034054 |
| SARS-CoV-2/human/CHN/CN3/2020 | GenBank | MT407652 |
| SARS-CoV-2/human/CHN/Shanghai_CH-02/2020 | GenBank | MT627325 |
| SARS-CoV-2/human/CHN/WHUHnCoV011/2020 | GenBank | MT079851 |
| SARS-CoV-2/human/CHN/WHUHnCoV001/2020 | GenBank | MT079843 |
| SARS-CoV-2/human/CHN/WHUHnCoV004/2020 | GenBank | MT079846 |
| SARS-CoV-2/human/CHN/WHUHnCoV020/2020 | GenBank | MT079853 |
| SARS-CoV-2/human/CHN/WHUHnCoV002/2020 | GenBank | MT079844 |
| SARS-CoV-2/human/CHN/WHUHnCoV008/2020 | GenBank | MT079850 |

| | | |
|---|---|---|
| SARS-CoV-2/human/CHN/GZMU0047/2020 | GenBank | MT568640 |
| SARS-CoV-2/human/CHN/OS2/2020 | GenBank | MT407658 |
| SARS-CoV-2/human/CHN/Beijing_IME-BJ03/2020 | GenBank | MT291833 |
| SARS-CoV-2/human/CHN/231/2020 | GenBank | MT135042 |
| SARS-CoV-2/human/CHN/235/2020 | GenBank | MT135044 |
| SARS-CoV-2/human/CHN/Beijing_IME-BJ04/2020 | GenBank | MT291834 |
| SARS-CoV-2/human/CHN/WHUHnCoV005/2020 | GenBank | MT079847 |
| SARS-CoV-2/human/CHN/WHUHnCoV006/2020 | GenBank | MT079848 |
| Yunnan/IVDC-YN-003/2020 | GISAID | EPI_ISL_408480 |
| SARS-CoV-2/human/CHN/WHUHnCoV007/2020 | GenBank | MT079849 |
| SARS-CoV-2/human/CHN/GZMU0044/2020 | GenBank | MT568639 |
| SARS-CoV-2/human/CHN/WHUHnCoV007/2020 | GenBank | MT079849 |
| SARS-CoV-2/human/CHN/GZMU0044/2020 | GenBank | MT568639 |
| SARS-CoV-2/human/CHN/GZMU0048/2020 | GenBank | MT568641 |
| Meizhou_MZ02/2020 | GenBank | MT510727 |
| Meizhou_MZ01/2020 | GenBank | MT510728 |
| SARS-CoV-2/human/CHN/GZMU0016/2020 | GenBank | MT568635 |
| SARS-CoV-2/human/CHN/WHUHnCoV012/2020 | GenBank | MT079852 |
| SARS-CoV-2/human/CHN/Shanghai_CH-03/2020 | GenBank | MT622319 |
| SARS-CoV-2/human/CHN/CN4/2020 | GenBank | MT407653 |
| RaTG13 | GenBank | MN996532.1 |
| RmYN02 | GISAID | EPI_ISL_412977 |

**Table S2.** Nucleotide frequencies and mutation rates used in the parametric simulations.

| Nucleotide | A | G | C | T |
|---|---|---|---|---|
| Frequency | 0.298101 | 0.183265 | 0.195801 | 0.322833 |

| Mutations | A→C | A→G | A→T | C→G | C→T | G→T |
|---|---|---|---|---|---|---|
| Rate | 0.767496 | 5.729515 | 1.093530 | 0.575406 | 16.142857 | 1 |

(((((((((((((SARS-CoV-2/human/CHN/Beijing$_I$ME-BJ01/2020:0.000069,Shenzhen/HKU-SZ-002/2020:0.000001):0.000001,USA/AZ1/2020:0.000034):0.000001,(Japan/TY-WK-012/2020:0.000034,(Japan/TY-WK-501/2020:0.000001,Japan/TY-WK-521/2020:0.000001):0.000001):0.000034):0.000001,USA/TX1/2020:0.000137):0.000001,Shenzhen/HKU-SZ-005/2020:0.000068):0.000034,SARS-CoV-2/human/CHN/WHUHnCoV003/2020:0.000103):0.000001,(((SARS-CoV-2/human/CHN/CN2/2020:0.000034,SARS-CoV-2/human/CHN/CN1/2020:0.000001):0.000001,SARS-CoV-2/human/CHN/CN5/2020:0.000034):0.000034,(((((SARS-CoV-2/IQTC01/human/2020/CHN:0.000001,Japan/KY-V-029/2020:0.000069):0.000068,((SARS-CoV-2/human/CHN/HZ-477/2020:0.000001,(((((((((SARS-CoV-2/human/CHN/HZ-178/2020:0.000001,(BetaCoV/Wuhan/YS8011/2020:0.000001,((((SARS-CoV-2/human/CHN/HZ-62/2020:0.000001,(((SARS-CoV-2/human/CHN/HZ-90/2020:0.000001,(SARS-CoV-2/human/CHN/HZ-48/2020:0.000001,Wuhan/WIV05/2019:0.000068):0.000001):0.000001,(SARS-CoV-2/human/CHN/Wuhan$_I$ME-WH05/2019:0.000001,SARS-CoV-2/human/CHN/HZ-49/2020:0.000001):0.000001):0.000001,(SARS-CoV-2/human/CHN/HZ-185/2020:0.000001,(((Wuhan/IPBCAMS-WH-02/2019:0.000001,SARS-CoV-2/human/CHN/HZ-79/2020:0.000001):0.000001,(SARS-CoV-2/human/CHN/GZMU0014/2020:0.000102,SARS-CoV-2/human/CHN/YN-0306-466/2020:0.000068):0.000001):0.000001,(SARS-CoV-2/human/CHN/Beijing$_I$ME-BJ05/2020:0.000034,(((SARS-CoV-2/IQTC02/human/2020/CHN:0.000034,(SARS-CoV-2/IQTC03/human/2020/CHN:0.000034,SARS-CoV-2/human/Guangzhou/IQTC05/2020:0.000034):0.000034):0.000069,USA/WI1/2020:0.000001):0.000001,SARS-CoV-2/human/CHN/Wuhan$_Y$B012506/2020:0.000001):0.000034):0.000001):0.000001):0.000001):0.000001):0.000001,hCoV-19/Thailand/61/2020:0.000001):0.000001,Wuhan/WH01/2019:0.000069):0.000001,((SARS-CoV-2/human/CHN/Wuhan$_Y$B012602/2020:0.000001,(SARS-CoV-2/human/CHN/Wuhan$_Y$B012504/2020:0.000001,SARS-CoV-

2/human/CHN/Wuhan_YB012605/2020:0.000001):0.000001):0.000001,SARS-CoV-
2/human/CHN/Wuhan_YB012611/2020:0.000103):0.000034):0.000001):0.000001):0.000001,SARS-
CoV-2/human/CHN/HZ-481/2020:0.000001):0.000001,China/WH-09/2020:0.000001):
0.000001,(((SARS-CoV-2/human/CHN/HZ-91/2020:0.000001,(((Wuhan/WH03/2020:
0.000001,SARS-CoV-2/human/CHN/HZ-576/2020:0.000001):0.000001,Taiwan/NTU02/2020:
0.000068):0.000001,(BetaCoV/Wuhan/WH19005/2019:0.000068,((SARS-CoV-
2/human/CHN/HZ-638/2020:0.000001,SARS-CoV-2/human/CHN/HZ-
551/2020:0.000001):0.000001,(SARS-CoV-2/human/CHN/WHUHnCoV021/2020:
0.000068,Wuhan/IPBCAMS-WH-05/2020:0.000034):0.000001):0.000001):0.000001):
0.000001):0.000001,(((BetaCoV/Japan/AI/I-004/2020:0.000069,SARS-CoV-
2/human/CHN/SH01/2020:0.000068):0.000001,((Wuhan/IPBCAMS-WH-04/2019:
0.000001,Wuhan/WIV06/2019:0.000001):0.000001,USA/CA9/2020:0.000034):0.000001):
0.000001,((((((SARS-CoV-2/human/CHN/HZ-162/2020:0.000001,((SARS-CoV-
2/human/CHN/Changzhou_JS27/2020:0.000001,SARS-CoV-2/Hu/DP/Kng/19-
027:0.000034):0.000001,SARS-CoV-2/Hu/DP/Kng/19-020:0.000001):0.000034):
0.000001,Wuhan/WIV04/2019:0.000001):0.000001,(hCoV-19/Germany/BavPat1/2020:
0.000001,(hCoV-19/Guangzhou/GZMU0037/2020:0.000001,(((SARS-CoV-
2/human/CHN/OS6/2020:0.000069,SARS-CoV-2/human/CHN/OS3/2020:
0.000034):0.000034,SARS-CoV-2/human/CHN/OS4/2020:0.000137):0.000001,hCoV-
19/Zhejiang/HZ103/2020:0.000034):0.000034):0.000001):0.000068):0.000001,Wuhan/IPBCAMS-
WH-03/2019:0.000034):0.000001,BetaCoV/Wuhan/WH19004/2020:0.000068):
0.000001,((((SARS-CoV-2/human/CHN/Fuyang_FY002/2020:0.000103,((SARS-CoV-
2/human/CHN/OS5/2020:0.000069,Brazil/SPBR-02/2020:0.000034):0.000034,SARS-
CoV-2/human/CHN/OS1/2020:0.000001):0.000001):0.000034,(SouthKorea/SNU01/2020:
0.000274,(Sweden/01/2020:0.000206,USA/CA2/2020:0.000034):0.000001):0.000001):
0.000001,Australia/VIC01/2020:0.000068):0.000034,USA/CA6/2020:0.000001):0.000001):
0.000001):0.000001):0.000001,Wuhan/IPBCAMS-WH-01/2019:0.000103):0.000001):
0.000001,2019-nCoV_WHU01:0.000001):0.000001,SARS-CoV-2/human/CHN/HZ-60/2020:
0.000001):0.000001,Hangzhou/HZ-1/2020:0.000001):0.000001,(Wuhan/WIV07/2019:
0.000068,(USA/CA5/2020:0.000068,USA/CA3/2020:0.000103):0.000001):0.000001):
0.000001,(((Wuhan/WIV02/2019:0.000034,BetaCoV/Wuhan/WH19008/2019:0.000001):
0.000001,USA/CA8/2020:0.000001):0.000034,Wuhan-Hu-1:0.000001):0.000001):

$$0.000001):0.000001,(SARS-CoV-2/human/CHN/Wuhan_OS52/2020:0.000137,(2019-nCoV_WHU02:0.000001,(Nepal/61/2020:0.000034,SARS-CoV-2/human/CHN/Beijing-01/2020:0.000034):0.000001):0.000001):0.000001):0.000001):0.000001,(SARS-CoV-2/human/CHN/CN3/2020:0.000034,SARS-CoV-2/human/CHN/Shanghai_CH-02/2020:0.000103):0.000034):0.000068,SARS-CoV-2/human/CHN/WHUHnCoV011/2020:0.000001):0.000001,((SARS-CoV-2/human/CHN/WHUHnCoV001/2020:0.000001,SARS-CoV-2/human/CHN/WHUHnCoV004/2020:0.000001):0.000001,((SARS-CoV-2/human/CHN/WHUHnCoV020/2020:0.000001,SARS-CoV-2/human/CHN/WHUHnCoV002/2020:0.000001):0.000001,SARS-CoV-2/human/CHN/WHUHnCoV008/2020:0.000001):0.000001):0.000068):0.000001):0.000001):0.000001,((SARS-CoV-2/human/CHN/GZMU0047/2020:0.000001,SARS-CoV-2/human/CHN/OS2/2020:0.000001):0.000001,(((SARS-CoV-2/human/CHN/Beijing_IME-BJ03/2020:0.000001,((SARS-CoV-2/105/human/2020/CHN:0.000001,SARS-CoV-2/human/CHN/231/2020:0.000001):0.000001,((SARS-CoV-2/233/human/2020/CHN:0.000034,SARS-CoV-2/human/CHN/235/2020:0.000001):0.000001,SARS-CoV-2/human/CHN/Beijing_IME-BJ04/2020:0.000034):0.000001):0.000001):0.000001,SARS-CoV-2/human/CHN/HN03/2020:0.000034):0.000068,SARS-CoV-2/human/CHN/Beijing_IME-BJ07/2020:0.000171):0.000001):0.000001):0.000001,((SARS-CoV-2/human/CHN/WHUHnCoV005/2020:0.000034,(USA/IL2/2020:0.000069,USA/CA1/2020:0.000069):0.000103):0.000001,(SARS-CoV-2/human/CHN/WHUHnCoV006/2020:0.000001,((((SARS-CoV-2/human/CHN/Yunnan-01/2020:0.000001,Yunnan/IVDC-YN-003/2020:0.000001):0.000069,(SARS-CoV-2/human/CHN/WHUHnCoV007/2020:0.000034,((SARS-CoV-2/human/CHN/GZMU0044/2020:0.000001,(SARS-CoV-2/IQTC04/human/2020/CHN:0.000068,USA/CA7/2020:0.000034):0.000001):0.000001,Taiwan/NTU01/2020:0.000001):0.000001):0.000001):0.000001,SARS-CoV-2/human/CHN/GZMU0048/2020:0.000001):0.000001,((Wuhan/WH04/2020:0.000001,((Meizhou_MZ02/2020:0.000069,Meizhou_MZ01/2020:0.000034):0.000068,SARS-CoV-2/human/CHN/GZMU0016/2020:0.000001):0.000001):0.000001,SARS-CoV-2/human/CHN/WHUHnCoV012/2020:0.000001):0.000001):0.000001):0.000001):0.000001):0.000034,SARS-CoV-2/human/CHN/Shanghai_CH-03/2020:0.000068):0.000001,SARS-CoV-2/WA2/human/2020/USA:0.000103):0.000001,USA/WA1/2020:0.000001):0.000001,SARS-CoV-$$

---

$2/human/CHN/CN4/2020:0.000137):0.007195;$

### References


Baele, G., Lemey, P., and Suchard, M. A. 2016. Genealogical working distributions for bayesian model testing with phylogenetic uncertainty. *Systematic Biology*, 65(2): 250–264.

Spielman, S. J. and Wilke, C. O. 2015. Pyvolve: a flexible python module for simulating sequences along phylogenies. *PloS one*, 10(9): e0139047.
